# Cardiac RyR N-terminal region biosensors for FRET-based high-throughput screening

**DOI:** 10.1101/2021.02.07.430153

**Authors:** Jingyan Zhang, Siobhan M. Wong King Yuen, Jacob A. Schwarz, Levy M. Treinen, Ching-Chieh Tung, Robyn T. Rebbeck, Kaja Berg, Bengt Svensson, Courtney C. Aldrich, David D. Thomas, Filip Van Petegem, Razvan L. Cornea

**Affiliations:** Department of Biochemistry, Molecular Biology and Biophysics, University of Minnesota, Minneapolis, MN 55455, USA; Department of Biochemistry and Molecular Biology, Life Sciences Institute, University of British Columbia,V6T 1Z3 Vancouver, BC, Canada; Department of Medicinal Chemistry, University of Minnesota, Minneapolis, MN, 55455, USA

**Keywords:** Ryanodine receptor, N-terminal region, FRET, fluorescence lifetime, high-throughput screening, myopathy

## Abstract

The N-terminal region (NTR) of the ryanodine receptor (RyR) calcium channels is critical to the regulation of Ca^2+^ release during excitation-contraction coupling. NTR hosts numerous mutations linked to skeletal and cardiac myopathies (RyR1 and RyR2, respectively), highlighting its potential as therapeutic target. Here, we labeled the NTR of mouse RyR2 at subdomains A, B, and C with donor and acceptor pairs for fluorescence resonance energy transfer (FRET), obtaining two biosensors. Using fluorescence lifetime (FLT)-detection of intramolecular FRET, we developed high-throughput screening (HTS) assays with the biosensors to identify small-molecule modulators of RyR. We screened a 1280-compound validation library and identified several hits. Hits with saturable FRET dose-response profiles, and previously unreported effects on RyR activity, were further tested using [^3^H]ryanodine binding to isolated sarcoplasmic reticulum vesicles, to measure their effects on full-length RyR opening in its natural membrane environment. We identified three novel inhibitors of both RyR1 and RyR2, and two RyR1-selective inhibitors at nanomolar Ca^2+^. These compounds may function as inhibitors of leaky RyRs in muscle. Two of these hits activated RyR1 only at micromolar Ca^2+^, highlighting them as potential activators of excitation-contraction coupling. These results indicate that large-scale HTS using this platform can lead to compounds with potential for therapeutic development.

## Introduction

The ryanodine receptor (RyR), a homotetrameric (~2.2 MDa) channel embedded in the sarcoplasmic reticulum (SR) membrane, is responsible for Ca^2+^ release from SR storage to cytosol, to enable excitation-contraction coupling(1,2). Defective RyR in skeletal (RyR1) or cardiac (RyR2) muscle cells perturbs Ca^2+^ release behavior, which can result in acquired and inherited myopathies (3–9). Therefore, RyRs are potential therapeutic targets for treating skeletal and cardiac myopathies, arrhythmias, and heart failure (HF) (9–13). Physiologically, the opening and closing of RyR channels are tightly regulated by small molecules, ions, and proteins(14), many of them binding to the enormous RyR cytoplasmic portion, with functional effects allosterically transduced to the cytoplasmic pore through long-range domain-domain interactions (15). The understanding of regulation by these domains has been greatly advanced by recent high-resolution cryo-electron microscopy (cryo-EM) RyR structures (15–19). The complex and rich regulation mechanisms of RyR domains offer opportunities to discover therapeutics that act either directly on RyR itself or modulate interacting proteins that determine RyR function.

Aiming to discover RyR regulators for therapeutic purposes, our group has previously developed a high-throughput screening (HTS) platform based on fluorescence-labeled RyR regulators FK506 binding protein 12.6 (FKBP) and calmodulin (CaM) (20,21). In that work, fluorescence lifetime (FLT) detection of FRET between the two labeled regulators was used as an assay to identify small-molecule modulators of RyR1. Using this assay, compounds have been identified to decrease RyR1 Ca^2+^ leak in skeletal muscle SR membrane vesicles and mechanically skinned muscle fibers (20,21), and were further shown to mitigate force loss in a muscular dystrophy mouse model (22). However, the binding sites of these molecules are difficult to identify within the full-length RyR, which limits progress on understanding their mechanisms of action and hinders structure-activity relationship studies. Similar limitations reside with other recently reported HTS platforms for discovery using full-length of RyRs (23). An attractive solution to this problem would be to develop an HTS platform based on constructs with fluorescent proteins directly attached to the RyR (24). However, that is currently quite a challenging proposition for HTS applications, primarily due to the inadequate level of expression of the biosensor constructs. As a practical alternative, we are now exploring the use of an essential, crystallizable RyR fragment.

The N-terminal region (RyR residues 1-547; termed NTR) is important for the functional regulation of RyR channels (25). The regulatory role of the NTR is partially attributed to its structural location in the RyR tetrameric channel complex (17,26). Recent high-resolution cryo-EM structures of the tetrameric RyR1 and RyR2, and earlier docking of the NTR to lower-resolution cryo-EM RyR structures, reveal that each NTR is comprised of three domains, A, B, and C (hence the NTR is often termed the ABC domain.). These are located at the first tier of the RyR-channel core structure (facing the cytosol), where NTRs interact with each other through their respective domains A and B, forming a tetramer that delimits a vestibule leading to the RyR channel pore (Figure 1B) (18,27). The NTR also directly interfaces with the RyR bridging solenoid, and then couples with the other cytoplasmic domains to allosterically regulate the channel function (18). The structural evidence of this long-range allosteric coupling between the NTR and other RyR cytoplasmic domains correlates with extensive functional studies (28–31), to solidify the early “domain-zipping” hypothesis. This hypothesis postulates that in the normal resting state, a large N-terminal portion of RyR1 and RyR2 interacts with a stretch of a the central portion of the RyR sequence making close contact (“domain zipping”), while on physiological stimulation, these critical inter-domain contacts are weakened (“domain unzipping”) (25), probably becoming more dynamic, and thus promoting channel opening (32). Very recently, it was also proposed that NTR self-association is the “gatekeeper” of RyR2 channel activity. Specifically, a stable N-terminal tetramer maintains the RyR2 channel closure, whereas disruption of this tetramer results in channel dysfunction (33). In addition, the NTR is one of the three major clusters of mutations associated with skeletal and cardiac myopathies such as malignant hyperthermia (MH) and catecholaminergic polymorphic ventricular tachycardia (CPVT) (26,27,34–36). Many of these disease-associated mutations in the NTR are found at the interfaces between domains A, B, and C. Some of these mutations reduce packing and destabilize the domain folding of the NTR, resulting in differences in structure or physical properties compared to the wild-type (WT)-NTR. For example, CPVT-associated mutant R420Q-NTR lacks a Cl^−^ ion that has been observed at a central location of convergence among the three domains of WT-NTR. This Cl^−^ appears to confer RyR2 WT-NTR structural stability (35). Taken together, the NTR is a structurally and functionally important component of the RyR channels, and structural perturbation of the NTR has the potential to affect RyR channel activity.

**Figure 1.**
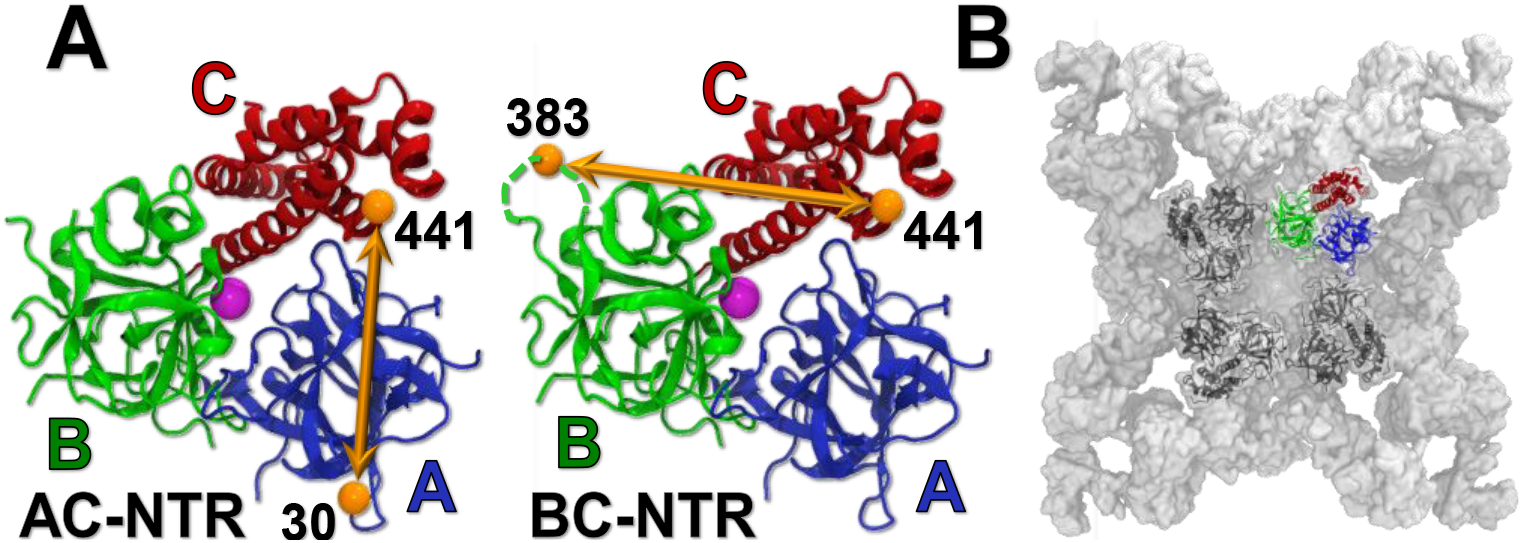
A) Two FRET biosensors, AC-NTR and BC-NTR, were engineered using the NTR of mouse RyR2. For FRET labeling, Cys residues were substituted at K30 (domain A), R383 (domain B), or K441 (domain C) shown in the crystal structure of residues 1-547 of RyR2 (PDB ID: 4L4H).^25^ Dashed lines represent the residues that are not resolved in the crystal structure. The purple sphere represents the chloride anion. B) Location of the NTR within the cryo-EM RyR2 structure (PDB ID: 5GO9).^17^

Given the critical role of the NTR in the regulation of RyR, we have engineered NTR constructs for use as FRET biosensors for an HTS platform to identify small molecules that regulate the activity of full-length RyR. We introduced donor-acceptor FRET pairs at appropriately distanced sites in different domains of the NTR (Figure 1A), then used an FLT plate-reader to measure changes in FLT-detected FRET efficiency as an indication of structural changes in the NTR due to small-molecule binding. Initial hit compounds obtained through FLT-FRET HTS of a small library were further validated using a dose-response format of the same FRET assay, and this was followed by functional assays (ryanodine binding measurements) using full-length RyR in SR membrane vesicles of skeletal and cardiac muscles to further confirm their effects on RyR channels.

## Results

### Engineering FRET biosensor constructs based on the NTR of mouse RyR2

To build a FRET biosensor using isolated NTR, seven solvent-exposed Cys residues were first mutated into Ala, to obtain a Cys-light NTR. Per construct, two of three residues, K30 in domain A, R383 in domain B, and K441 in domain C, were then mutated to Cys, to provide the thiol labeling sites. Based on the crystal structure shown Figure 1, these three Cys residues are fully exposed to the solvent and accessible for labeling. We hypothesized that FRET-detected changes in their relative positions could provide an assay reflecting structural changes that occur in the NTR in response to ligand binding. Each test-biosensor construct contains two of the introduced Cys residues, K30C/K441C (termed AC-NTR), and R383C/K441C (termed BC-NTR). The inter-probe distances in the AC- and the BC-NTR constructs were estimated at 42 Å and 50 Å, respectively (note that R383, and K441 are not resolved in the crystal structure PDB ID 4L4H; instead, the nearby residues were used to estimate the inter-probe distances; Figure1) (35). Based on the predicted distances between the labeling sites in the two constructs, Alexa Fluor-488 and Alexa Fluor-568 fluorescent dyes were chosen as FRET donor and acceptor, respectively. Both constructs were labeled with a ratio of 2:3 between the donor and acceptor, to maximize FRET efficiency (Eq. 1). Inter-probe distances of the two biosensors were determined using previously described methods of multi-exponential analysis of the FLT-detected FRET measurements (Figure S1) (37), and were consistent with the distances predicted based on the crystal structure.

### Evaluation of the NTR FRET-biosensor sensitivity to small-molecule ligands

The AC-NTR and BC-NTR FRET constructs were evaluated by comparing the change in the distances of WT- and R420Q-NTR determined via FRET data analysis in the presence of different concentrations of Cl^−^. The crystal structure of WT-NTR contains a Cl^−^ ion at a central position, stabilizing the protein through electrostatic interactions with nearby residues of R420, R298, and R276 (35). In practice, millimolar [Cl^−^] was necessary to stabilize NTR in solutions during the purification. However, Cl^−^ was not observed in the crystal structure of the R420Q-NTR mutant. We probed the reported structural difference between WT- and R420Q-NTR, and their responses to Cl^−^, to determine if the FRET assay of the two constructs (AC and BC) is sensitive to small ligands. Multiexponential analysis of the FLT-detected FRET measurements was performed as previously described (38–41). A single-Gaussian model, described by the inter-probe distance (R) and its full-width at half-maximum (FWHM), provided the best fit of the WT AC and BC, and R420Q-BC FRET data sets, such as those illustratively shown in Figure S1. A summary of the FRET data fitting parameters is shown in Table S1, and Figure 2 illustrates R changes for these constructs in the presence of high (100 mM) and low (1 μM) Cl^−^. We observe ~10 Å longer inter-probe distances within WT-NTR with both FRET-labeled constructs, in high [Cl^−^] vs. low [Cl^−^] buffer, indicating that the two sensors can report ligand-induced changes in the NTR structure Figure 2, left). However, we observed no significant change in the inter-probe distances of the R420Q mutant, in high vs. low [Cl^−^] buffers (Figure 2, right), indicating that the R420-NTR construct is insensitive to Cl^−^. This is consistent with Cl^−^ not being a ligand for R420Q (35). FWHMs of the R distributions were quite similar under both [Cl^−^] conditions, suggesting similar levels of disorder under the different [Cl^−^] conditions (Table S1). This result indicates that the tested WT-NTR FRET constructs could be used as biosensors in studies to resolve small-molecule ligands of the NTR, such as done via HTS of chemical libraries containing drug-like molecules.

**Figure 2.**
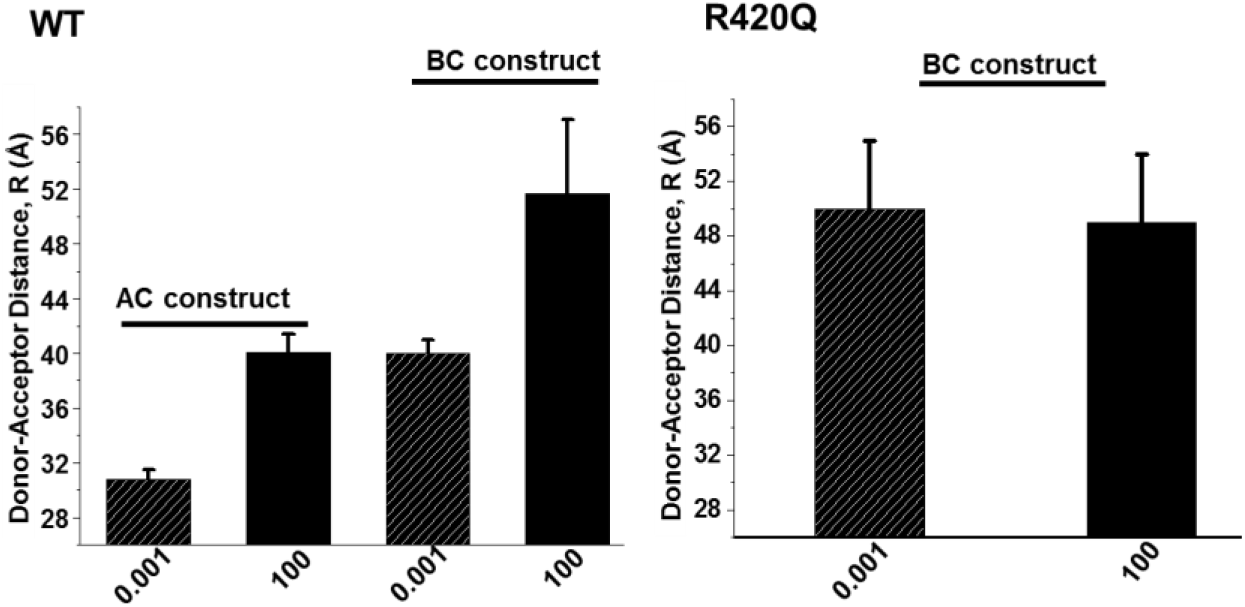
Summary of inter-probe (donor-acceptor) distances determined from analysis of FRET assays of WT- and R420Q NTR, measured using AC- and BC-NTR constructs in the presence of low (0.001 mM) and high (100 mM) Cl^−^ in the buffer, respectively (based on fitting parameters shown in Table S1 from the analysis of data such as shown in Figure S2). Results are shown as mean ± SD (n = 3).

### FRET-based HTS using biosensor constructs derived from RyR2 NTR

FRET-labeled AC- and BC-NTR constructs were used to screen the 1280-compound LOPAC collection, which is a typical step for biosensor validation. HTS was carried out in 1536-well black-wall/black-bottom plates, formatted as described in our previous work (21,42). A donor-only labeled construct was included for each screen as a control used to filter out the compounds that have a fluorescence signal overlapping with the donor fluorescence spectrum, and which might therefore interfere with the FLT reading. Both the FLT waveforms and fluorescence spectra of each well in each plate were acquired after 20, 60, and 120 min of incubation with the LOPAC molecules. The FLT change of each well in the plate in the presence of LOPAC molecules was used as a high-precision assay of a structural change in the biosensor due to binding of a compound, previously shown to be ~30-fold more precise than typically possible using fluorescence intensity (20). For a quantitative evaluation of the biosensors, we used the coefficient of variation (CV% = 100 x SD/means), which was 0.88±0.13% for the AC construct and 1.41±0.19% for the BC construct. Based on these CV values, a reference compound (positive control) would have to produce ΔFLT of at least 0.35 ns or 0.53 ns in order to achieve the “excellent” rating (i.e., Z’≥0.5) for an assay driven by the AC or BC construct, respectively (per Eq. (2)) (43).

Library compounds that changed FLT beyond 5SD from the means of the DMSO controls were selected as hits. False positive hits were filtered out based on changes in donor-only FLT, ratio between channel 1 and channel 2 intensity, fluorescence peak intensity and shape, and similarity index (44). Figure 3 (left panel) illustrates a representative outcome of the HTS. Each data point corresponds to the ΔFLT caused by one compound from LOPAC. Table S2 summarizes the Hit numbers and reproducibility at 20, 60, and 120 minutes incubation with the WT-AC and -BC constructs. Hit identities are indicated in Figure 3 (right panel). For each construct, these hits were found in at least two of the three runs of the screen. The screens using the AC-NTR construct picked up more hits than BC-NTR, as shown in Table S2. Longer incubation times resulted in more hits, which may be due to chemical reaction between the biosensor and drug molecules (Table S2 and Figure S2), or to aggregation of the NTR caused by the presence of drug molecules, or instability of the protein itself. All hits previously known to promote protein aggregation, as found in the Aggregator Advisor database (45), were not pursued in our next steps. Most of the remaining 18 hits are previously unreported RyR ligands (Figure 3, right panel).

**Figure 3.**
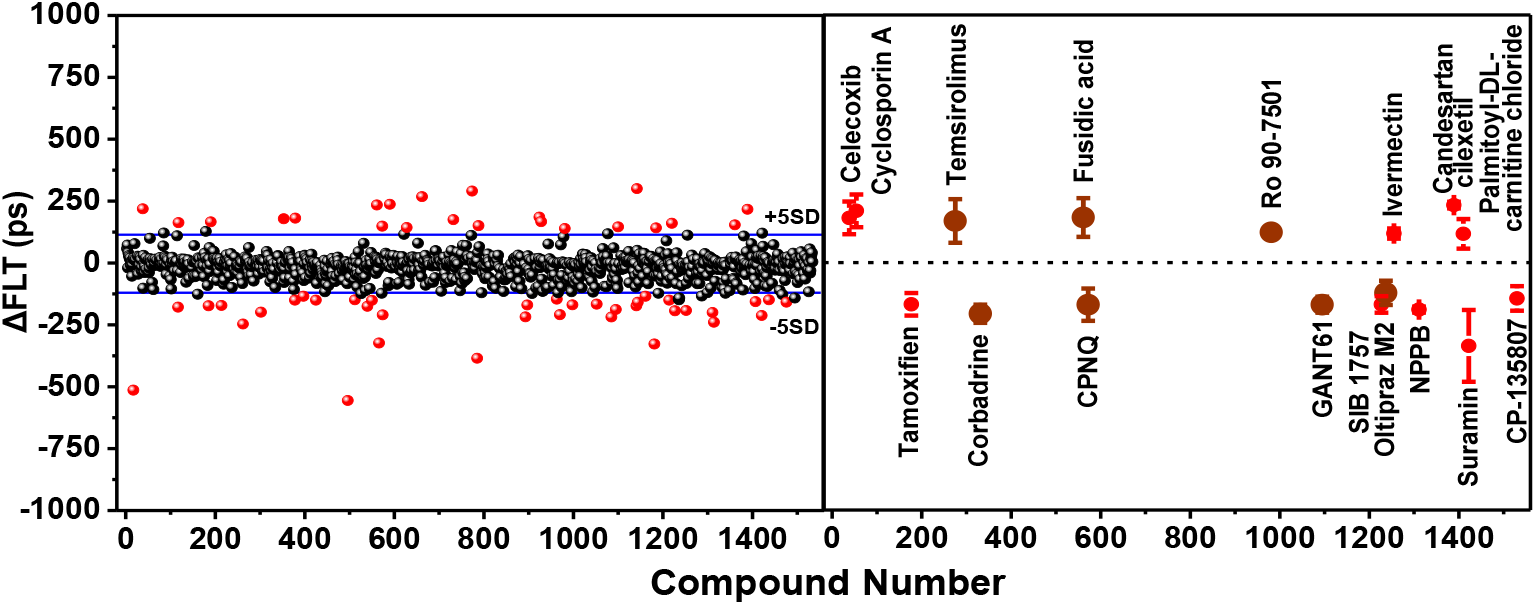
Left: A representative FLT-based FRET HTS run with the labeled AC-NTR construct. Dotted lines indicated the 5SD threshold. Right: 18 reproducible hits (i.e., identified in ≥2 of 3 HTS runs) from AC- or BC-NTR constructs. These were further assessed using FRET dose-response measurements (Figure 4). Dark red symbols indicate the hits selected for further testing. Right panel results are shown as mean ± SD (n = 3).

### FRET dose-response of the initial hits

To confirm the 18 initial hits, their FRET dose-responses were measured using the same method as in the primary HTS. In a range of 0-100 μM concentrations, the hits that decreased FLT in HTS also decreased FLT in this more detailed retest, and the same agreement was true for the Hits that increased FLT. Compounds tamoxifen and CP-31398 decreased FLT of the donor-only sample by > 50% of the FLT change in the donor-acceptor sample, and therefore were designated interfering compounds and were excluded from further studies. Hits that showed ΔFLT < 2% in these dose-response studies were also excluded from further evaluation. Based on these criteria, we narrowed down the number of hits for further evaluation to the eight compounds indicated by dark red symbols in Figure 3 (right panel). These are CPNQ, oltipraz M2, GANT61, corbadrine, fusidic acid, tacrolimus, temsirolimus, and Ro 90-7501 (chemical names shown in Table S3).

Individual FRET dose-response curves of the eight remaining hits, along with their chemical structures are shown in Figure 4. The solid lines represent fits of the experimental data using the Hill function. Among the hits that decreased FLT (Figure 4a), CPNQ had the strongest effect (ΔFLT ≈ 0.4 ns), which was similar to corbadrine, albeit the latter had lower potency (~ 3-fold higher EC_50_). Both GANT61 and oltipraz M2 decreased FLT but did not reach saturation within the tested concentration range, indicating a lower potency compared to corbadrine and CPNQ. Among the hits that increased FLT (Figure 4b), fusidic acid and tacrolimus exhibited a stronger effect than the other two compounds in this group (ΔFLT ≈ 0.25 ns). Temsirolimus and Ro 90-7501 produced smaller increases in FLT, but with a higher potency (EC_50_ ~0.1 μM). Temsirolimus and the related tacrolimus (aka FK506) are known ligands of FKBP, and they belong to a group of compounds thought to indirectly influence the RyR function, by preventing FKBP-RyR binding (46). Our results (Figure 4b) indicate that these compounds may also directly interact with the RyRs, a hypothesis to be explored in future studies. Tacrolimus and Ro 90-7501 had already been identified as RyR regulators in our previous HTS campaigns (via a FRET HTS assay that uses full-length RyRs), and their RyR activatory and inhibitory effect (respectively) have been reported (20,21). The remaining five compounds are previously unknown RyRs ligands, and we further evaluated them by [^3^H]ryanodine binding assays (shown below).

**Figure 4.**
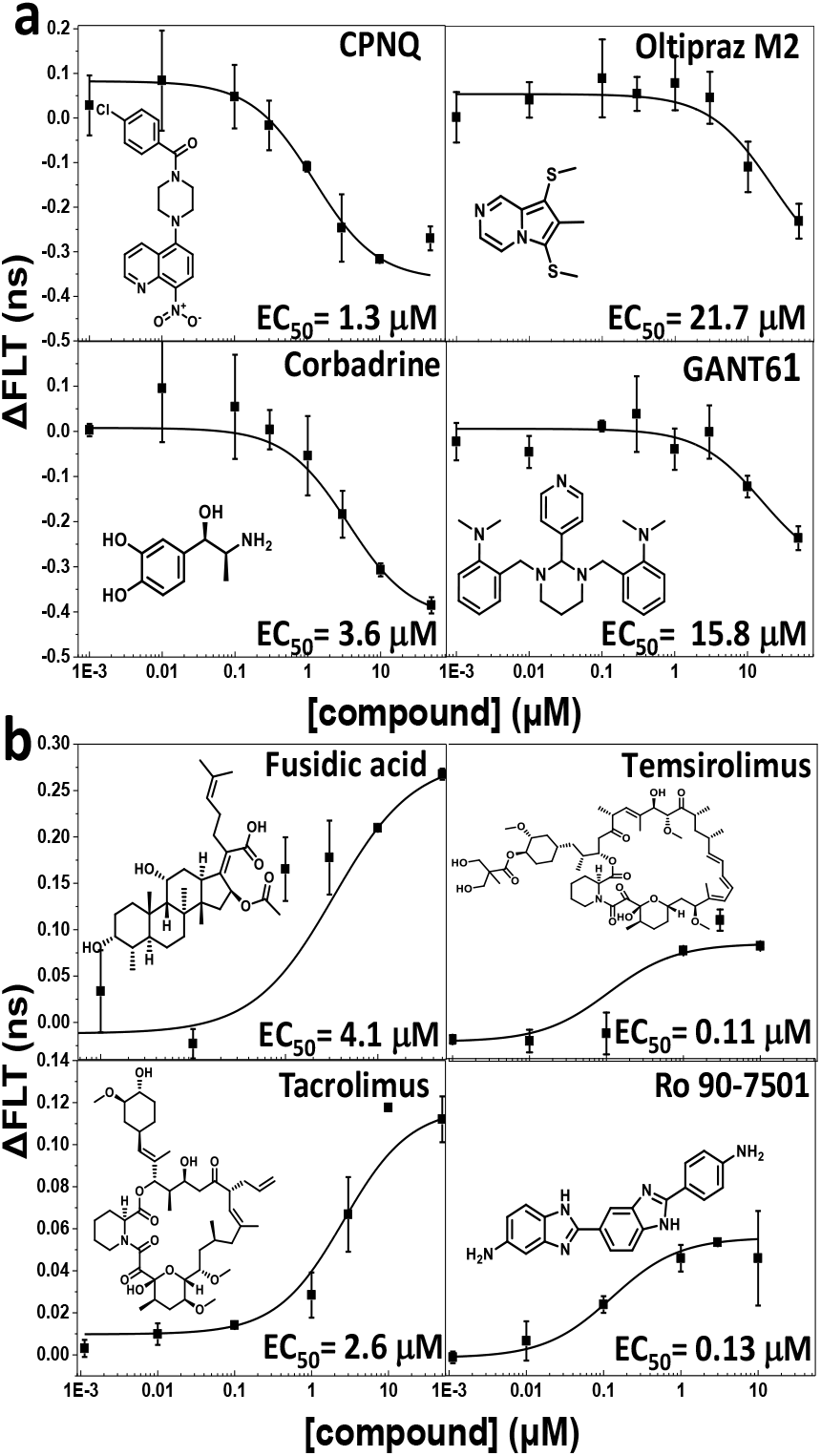
FRET dose-response curves of the hits measured by the AC-NTR biosensor. Compounds that decreased (a) and increased lifetime (b) of the FRET biosensor. Plots are shown relative to the values for no-drug control (DMSO), mean ± SD (n = 3).

### Effect of hits on [^3^H]ryanodine binding to full-length RyR1 and RyR2

To functionally evaluate the hits that had a clear FRET dose-response profile and previously unreported effects on RyR function, we performed [^3^H]ryanodine binding assays using SR membrane vesicles isolated from pig skeletal and cardiac muscle. This is a well-established method that is often used to determine the activity of RyR channel in their native membrane environment (47), and is therefore routinely used to evaluate hit effects on RyRs (20,21,23). The increase or decrease of bound [^3^H]ryanodine correlates with the fractional population of RyR open channels. [^3^H]ryanodine binding assays were carried out in the presence of low Ca^2+^ (30 nM) and high Ca^2+^ (30 μM), corresponding to the resting and contracting muscle conditions, respectively. This is to glean early insight in the potential effect of a hit on the resting RyR leak, as well as on RyR activation during excitation-contraction coupling.

RyR1 and RyR2 share more than 65% sequence similarity, and the NTRs of RyR1 and RyR2 also highly resemble each other in terms of sequence and structure (48). Thus, modulators of one isoform might also affect the other. Therefore, we performed [^3^H]ryanodine binding analyses to determine the dose-dependent effects of the novel FRET hits (from Figure 4) on RyR1 and RyR2 function, in skeletal (Figure 5, left) and cardiac membrane vesicles (Figure 5, right), respectively.

**Figure 5.**
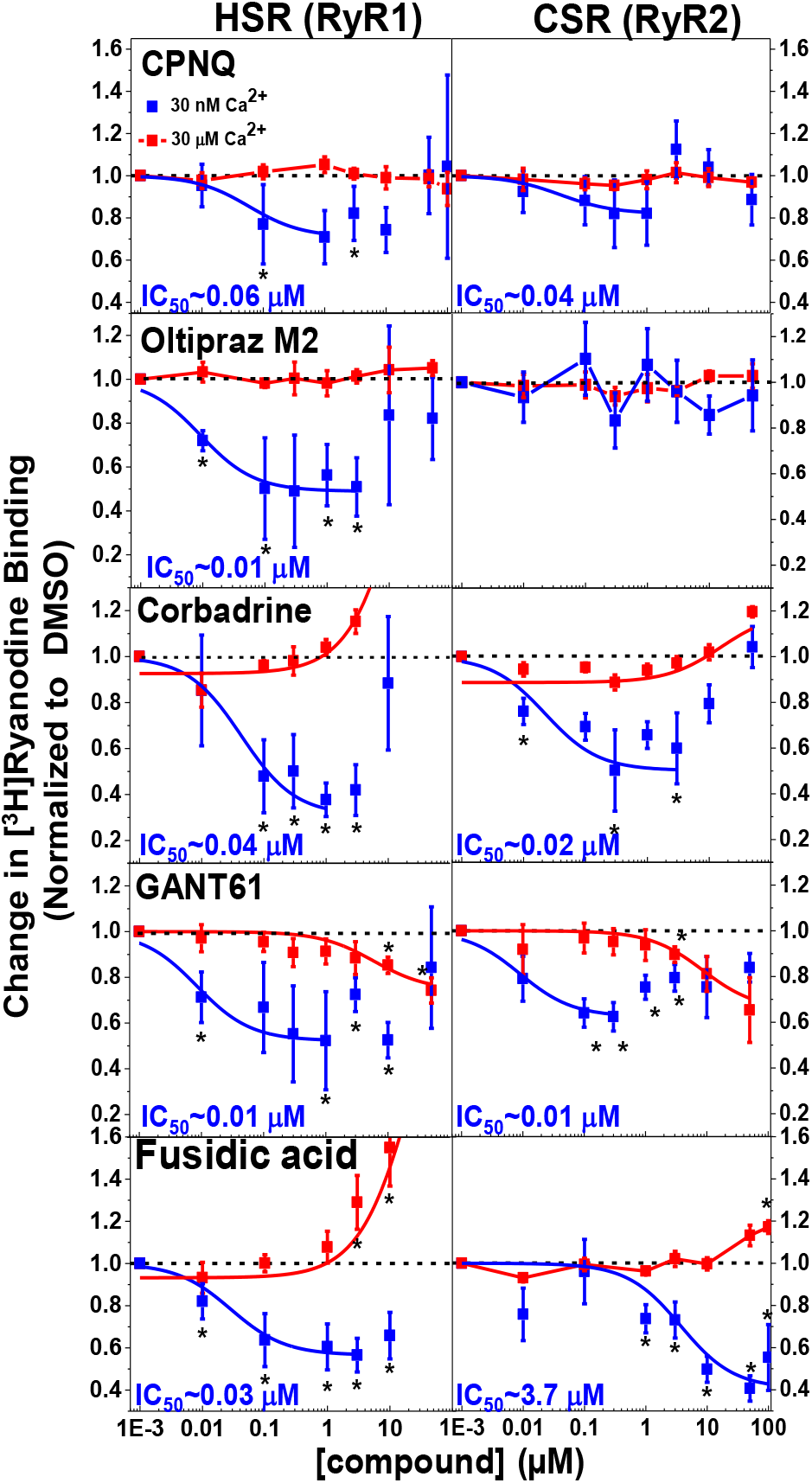
Dose-dependent effects of the hits on [^3^H]ryanodine binding to skeletal SR (HSR, left) and cardiac SR (CSR, right) membranes at 30 nM (blue) and 30 μM (red) free Ca^2+^. Results are shown normalized relative to the values for no-drug control (DMSO), respectively, mean ± SE, n = 4. Data were fit using the Hill function.*Significantly different from control using unpaired Student’s t-test, p > 0.05.

For skeletal SR (RyR1) at 30 nM Ca^2+^ (Figure 5, left panels, blue curves), all tested hits inhibited [^3^H]ryanodine binding in a concentration-dependent manner, with nanomolar IC_50_ values (as indicated). All compounds showed biphasic responses, with inhibition by submicromolar hit gradually turning into activation at the highest concentrations tested (Figure 5, left; Figure S3). Maximum inhibitory effects observed range from approximately 30% for CPNQ, to 50% for oltipraz M2, GANT61, and fusidic acid, to 70% for corbadrine (Figure 5, left; Figure S3).

For skeletal SR at 30 μM Ca^2+^ (Figure 5, left panel, red curves), both CPNQ and oltipraz M2 only slightly affected the [^3^H]ryanodine binding to RyR1 throughout the tested concentration range, and GANT61 showed a gradual inhibition at concentrations >1 μM. However, concentrations >1 μM of corbadrine and fusidic acid strongly increased [^3^H]ryanodine binding.

For cardiac SR (RyR2) at 30 nM Ca^2+^ (Figure 5, right panel, blue curves), the pattern of inhibitory effects by four of the hits was largely similar to RyR1. Oltipraz M2 was the exception, as it had almost no effect on RyR2. Inhibition of RyR2 by fusidic acid had micromolar IC_50_ (3.7 μM), vs. nanomolar (0.03 μM) for RyR1.

For cardiac SR at 30 μM Ca^2+^ (Figure 5, right panel, red curves), CPNQ and oltipraz M2 did not significantly alter [^3^H]ryanodine binding, similar to HSR. Corbadrine and fusidic acid showed slight activation at high concentrations, but much smaller than with HSR. GANT61 was an inhibitor at concentration >1 μM, similar to HSR.

## Discussion

RyR channels are under intense study as potential therapeutic targets for diseases ranging from rare ryanopathies to high-prevalence degenerative syndromes. Here, we used the FRET-labeled NTR to detect compound binding, as this is a domain that is critically involved in RyR function. Specifically, NTR assembles into a tetrameric vestibule (Figure 1B) that is sterically coupled to the channel pore regulation. The NTR is also one of the “hot-spots” of disease-associated mutations of RyR (26,27,34,35). Structural perturbation of the NTR itself or of its interaction with adjacent domains leads to disruption to the open/closed properties of RyR channels. In the related IP3 Receptor, the corresponding NTR forms the binding site for the activating ligand IP3 (7,49), which alters the conformation of the NTR. Therefore, small-molecule ligands of the NTR in RyRs have high potential to functionally regulate the full-length RyR channels, and be developed to obtain therapeutic agents against RyR dysfunction.

To exploit the unique structural features and functional importance of the NTR, and the well-established high-precision FLT detection of FRET, we constructed two NTR biosensors by substituting Cys at two residues located in A and C, or B and C sub-domains of the Cys-light NTR, and labeled these Cys pairs with fluorescent probes suitable as FRET donor and acceptor (Figure 1). The two NTR biosensors with the fluorescent probes located at different domains are expected to be sensitive to different structural changes and, therefore, detect different modulators of the NTR. Indeed, different inter-probe distances are reported by the two biosensors in response to chloride ions, validating that the constructed biosensors are responsive to changes in NTR structure/conformation (Figure 2), and suggesting that these constructs could be used in FRET-based HTS assays for RyR-targeted drug discovery. These results also support the hypothesis that binding or unbinding of chloride to the RyR2 NTR may regulate channel activity (35).

FRET-based HTS of the LOPAC with these two biosensor constructs identified hits that either increased or decreased the FLT of the biosensors (Figure 3). Screening with AC-NTR picked up more hits than BC-NTR, possibly due to the probe at K30 in the A domain being more sensitive than R383 in the B domain. Also, more hits decreased FLT than increased it (Figure S2). Based on reproducibility and chemical and physical properties, we selected 18 initial hits (Figure 3b) for further testing. The list of hits was further narrowed down to eight compounds based on FRET dose-response curves (Figure 4) and compound availability. Among the eight compounds, three were previously identified as potential RyR regulators, leaving five newly identified compounds for testing via assays of [^3^H]ryanodine binding to full-length RyR1 and RyR2 (Figure 5).

Five of the eight identified hits are structurally and functionally different from known RyR modulators, such as 1,4-benzothiazepine derivatives (aka rycals, e.g., JTV519 and S107), or dantrolene (50–52). These five hits are also structurally unrelated to the RyR small-molecule modulators discovered using biosensor systems using labeled FKBP and CaM (21). Of the five validated hits, four shortened the FLT (increased FRET), which suggests a shorter inter-probe distance within NTR; and one lengthened FLT (decreased FRET), which suggests a longer inter-probe distance. All five FLT hits also affected [^3^H]ryanodine binding to RyR1 and 2, which provides experimental support to the hypothesis that NTR ligands can regulate the functional status of RyR channels, as suggested by high-resolution structures of RyR channels and many functional studies (17,26,33).

Among the five hits, oltipraz M2 exhibited strong isoform specificity by inhibiting RyR1 only in nanomolar [Ca^2+^]. Oltipraz M2 is a metabolite of cancer chemopreventive agent oltipraz. It has been shown to exert cytoprotective effects *via* antioxidation mechanisms (53,54). CPNQ, targets huntingtin and α-synuclein to lower their pathological effects in cellular models of Huntington’s and Parkinson’s diseases, respectively (55), and we show here that this compound significantly inhibited only RyR1 (and an inhibitory trend on RyR2) at low Ca^2+^. GANT61, a hexahydropyrimidine derivative antitumor agent (56), is the only Hit that inhibited RyR1 and 2 channels at both low and high Ca^2+^.

Corbadrine and fusidic acid affected [^3^H]ryanodine binding to RyR1 and 2 most strongly among the tested compounds (Figure 5). Thus, the [^3^H]ryanodine binding profiles in the presence of corbadrine and fusidic acid – in nanomolar vs. micromolar Ca^2+^ – are particularly attractive for potential RyR1 therapeutics. However, corbadrine has potentially significant liabilities due to its adrenaline-analogous structure and the very high binding affinity for α2-adrenergic receptors, suppressing peripheral vasoconstriction (57). For this reason, corbadrine might be unsuitable for repurposing, despite its favorable effects on RyR1 and RyR2 in isolated SR membranes. On the other hand, fusidic acid has a good safety record and excellent pharmacokinetic profile, and good tissue distribution (58). Fusidic acid is a bacteriostatic antibiotic derived from the fungus *Fusidium coccineum* and is used as a topical medication to treat skin infections by inhibiting bacterial protein synthesis (59). Fusidic acid is rarely used as an antibiotic, in the US, because of drug resistance. However, it is FDA approved, well tolerated and therefore attractive as a repurposing candidate, as is often the case with antibiotics (58). Derivatives of fusidic acid have been synthesized by others (60), and they could be tested in future studies focused on establishing structure-activity relationships relevant to RyRs.

Among the previously identified RyR regulators (20,21), tacrolimus and temsirolimus are known FKBP ligands thought to prevent FKBP binding to RyRs. Tacrolimus was also identified by HTS using a measurement of endoplasmic reticulum Ca^2+^ in HEK293 cells (23). Given that tacrolimus and temsirolimus are known high-affinity ligands of FKBP, it was somewhat unexpected to identify them as ligands of our NTR biosensors. This observation suggests that they have the potential to also interact directly with full-length RyR (15,61).

In summary, we developed an early-stage RyR-targeted drug discovery method exploiting high-throughput FLT-detection of FRET within a RyR2 NTR construct labeled with donor and acceptor, to identify small-molecule modulators of RyR1 and RyR2. We identified several novel RyR1 and RyR2 effectors. Results from these assays suggest that none of these compounds have deleterious effects on RyR1 and RyR2, and some (e.g., oltipraz M2 and fusidic acid) should be further pursued through studies using cell and animal models of skeletal and cardiac myopathies. Further studies will be necessary to determine the mechanisms of action of these new effectors and pursue medicinal chemistry development toward a RyR-targeted drug. Future HTS screening of the NTR biosensor against a larger chemical library will accelerate the discovery of RyR-targeted therapeutics.

## Experimental procedures

### Expression, purification, and labeling of NTR

The NTR of mouse RyR2 (residues 1-547) was cloned and expressed as previously described (35). The DNA construct contains a 6×His tag, a maltose-binding protein (MBP) tag, and a tobacco etch virus (TEV) protease cleavage site. The seven native solvent-exposed cysteine residues were substituted with alanine residues, and two constructs were engineered with Cys substitutions at surface residues R383 and K441 (in the B and C domains, respectively; termed BC) and at K30 and K441C (in the A and C domains, respectively; termed AC) for labeling with spectroscopic probes such as appropriate donor-acceptor fluorophores for FRET (for the purpose of this report, these are considered the WT forms of AC and BC). A variant of the BC construct corresponding to the disease-associated mutant (R420Q) was also prepared. For this site-directed mutagenesis work, we used the QuikChange (Stratagene) and Phusion High-Fidelity PCR (Thermo Fisher Scientific) kits. The WT and R420Q AC and BC protein constructs were expressed in *Escherichia coli* Rosetta (DE3) pLacI cells at 18°C, and purified with a protocol adapted from previously published work (26). An amylose column was first applied after the cells were lysed by ultra-sonication. The protein in the elution was then cleaved using TEV protease. The cleaved crude protein went through the second amylose column to separate from the MBP, and then followed by a Superdex S-200 size exclusion column in a buffer consisting of 250 mM KCl, 10 mM HEPES, and 14 mM β-mercaptoethanol (pH 7.4). The NTR protein (3 μM) was then incubated in 10 mM HEPES, 250 mM KCl (pH 7.4, 2 h, 21°C) with the fluorescent probes Alexa Fluor 488 maleimide (AF488, donor; 40 μM) and Alexa Fluor 568 maleimide (AF568, acceptor; 60 μM). The Alexa dyes were added last to the reaction mixture, from 10 mM DMSO stock solutions. These conditions were experimentally optimized to maximize FRET efficiency. The labeling efficiency was determined by comparing the molar concentration of the bound dye, determined from UV-Vis absorbance, to the protein concentration, determined from gel densitometry. Essentially complete labeling was confirmed from electrospray ionization mass spectrometry. Circular dichroism spectra of the protein before and after labeling were recorded by JASCO J-815 (Japan), confirming similar folding.

### Chemical library handling, preparation of 1536-well assay plates and sample loading

The 1536-well assay plates were prepared as described in our previous reports (21). The LOPAC (Sigma-Aldrich, MO, USA) collection received in 96-well plates was reformatted into 384-well polypropylene intermediate plates (Greiner Bio-One, Kremsmünster, Austria) using a multichannel liquid handler, BioMek FX (Beckman Coulter, Miami, FL, USA), then transferred to 384-well Echo Qualified source-plates (Labcyte Inc, Sunnyvale, CA, USA). Assay plates were prepared by transferring 5 nL of the 10 mM compound stocks in columns 3-22 and 27-46, and 5 nL DMSO in columns 1-2, 23-26 and 47-48 from the source plates to 1536-well black polypropylene plates (Greiner), using an Echo 550 acoustic dispenser (Labcyte Inc.). These assay plates were then heat-sealed using a PlateLoc Thermal Microplate Sealer (Agilent Tech., Santa Clara, CA, USA) and stored at −20°C prior to usage. On screening days, these plates were thawed out, and were loaded with labeled NTR (5 μL, 20 nM final concentration) using a Multidrop Combi automated sample dispenser.

### FRET measurements

For cuvette-based determinations of NTR construct response to Cl^−^, time-correlated single-photon counting (TCSPC) measurement of fluorescence lifetime (FLT) was used. Time-resolved fluorescence decays were recorded after excitation with a 481-nm subnanosecond pulsed diode laser (LDH-P-C-485, PicoQuant, Berlin, Germany). Emitted light was selected using a 517 ± 10 nm filter (Semrock, Rochester, NY) and detected with a PMH-100 photomultiplier (Becker-Hickl, Berlin, Germany). FLT waveforms of the donor only and donor-acceptor-labeled samples were acquired in media containing 60 nM biosensor, 10 mM HEPES, 250 mM KCl, pH 7.4 at 21°C), and analyzed to determine FRET distance relationships, as described in our previous publications (38–41).

HTS measurements were conducted in a prototype top-read FLT plate reader (FLT-PR, Fluorescence Innovations, Inc., Minneapolis, MN), which reads each 1536-well plate in ~3 min (62,63). AF488 donor fluorescence was excited with a 473 nm microchip laser from Concepts Research Corporation (Belgium, WI), and the emission signal was split into two channels by a dichroic mirror and acquired with 517 ± 10 nm (channel 1) and 535 ± 7 nm band-pass filters (channel 2) (Semrock, Rochester, NY). This instrument uses a unique direct waveform recording technology that enables high-throughput FRET detection at high precision. Fluorescence spectra of each assay plate were also recorded with a SpectraMax Gemini EM plate reader (Molecular Devices). FLT waveforms for each well of 1536-well plates were fit based on a one-exponential decay function using least-squares minimization global-analysis software. The FRET efficiency (E) was determined as the fractional decrease of donor FLT (τ_D_), due to the presence of acceptor (τ_DA_), using the following equation:

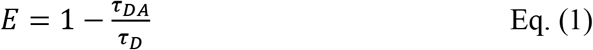

HTS assay quality was calculated based on the effect of positive control (10 μM compound) relative to negative control (0.1% DMSO) and measurement precision, as indexed by the Z’ factor (43):

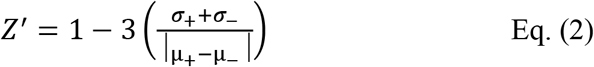

where σ_+_ and σ_−_ are the SDs of the positive and negative control τ_DA_, respectively; μ_+_ and μ_−_ are the means of the positive and negative control τ_DA_, respectively. A value of Z’≥ 0.5 indicates an excellent HTS assay, while 0.5>Z’≥0 indicates an adequate assay (43).

### Isolation of sarcoplasmic reticulum vesicles

Crude sarcoplasmic reticulum membrane vesicles were isolated from porcine longissimus dorsi muscle and porcine cardiac left ventricle tissue by differential centrifugation of homogenized tissue, as established previously (64). Skeletal heavy SR (HSR) vesicles, which are enriched with RyR1, were isolated by fractionation of crude skeletal SR vesicles using a discontinuous sucrose gradient, in accordance with our published work (64). All vesicles were flash-frozen and stored at −80°C.

### [^3^H] ryanodine binding to SR vesicles and data analysis

In 96-well plates, skeletal SR membranes (HSR, 1 mg/ml) and cardiac SR membranes (CSR, 3 mg/mL) were pre-incubated with 1% DMSO or hit compound for 30 min at 22°C in a solution containing 150 mM KCl, 5 mM GSH, 1 μg/mL Aprotinin/Leupeptin, 1 mM EGTA, and 65 μM or 1.02 mM CaCl_2_ (as determined by MaxChelator to yield 30 nM or 30 μM of free Ca^2+^, respectively), 0.1 mg/mL of BSA, and 20 mM K-PIPES (pH 7.0). Non-specific [^3^H]ryanodine binding to SR was assessed by addition of 40 μM non-radioactively labeled ryanodine. Maximal [^3^H]ryanodine binding was assessed by addition of 5 mM adenylyl-imidodiphosphate (AMP-PNP), and in the case of CSR, 20 mM caffeine was added. These control samples were each loaded over four wells per plate. Binding of [^3^H]ryanodine (7.5 and 10 nM for CSR and HSR, respectively) was determined following a 3 h incubation at 37°C and filtration through grade GF/B glass microfiber filters (Brandel Inc., Gaithersburg, MD, US) using a Brandel Harvester. In 4 mL of Ecolite scintillation cocktail (MP biomedicals, Solon, OH, USA), [^3^H]ryanodine retained on the filter was counted using a Beckman LS6000 scintillation counter Fullerton, CA).

### Statistical analysis

Errors are reported as the standard deviation of the mean (SD), except when noted. Statistical significance was determined by Student’s t-test one-way ANOVA followed by Tukey’s post-hoc test, as indicated, where p<0.05 was considered significant. EC_50_ values were derived from the fits to Hill equations.

## Data availability

All data are contained within the manuscript.

## Supporting information

Fluorescence lifetime waveforms and FRET distance distributions of the two biosensors in the presence of high and low chloride ions; HTS plots at different acquiring times; a full-size figure of the dose-dependent effect of corbadrine and fusidic acid on [^3^H]ryanodine binding to skeletal SR; summary of the distance and full-width at half maximum from the fitting of FLT-detected intramolecular FRET between donor-acceptor probes in the AC and BC biosensors corresponding to WT- and R420Q-NTR in the presence of high and low [Cl^−^], and hit rates obtained from 3 HTS runs are included in the supporting information.

## Acknowledgements

We thank Samantha Yuen for creating assay plates at the UMN Institute of Therapeutic Drug Discovery and Development. We thank Destiny Ziebol and Sarah Blakely for their assistance with manuscript preparation and submission. Spectroscopic measurements were performed at the UMN Biophysical Technology Center. This work was supported by NIH grants R01HL138539 (RLC) and R37AG026160 (DDT), an Individual Research Grant from the RYR-1 Foundation (RLC), and a Canadian Institutes of Health Research Grant 125893 (FVP). The content is solely the responsibility of the authors and does not necessarily represent the official views of the National Institutes of Health.

## Supporting information

**Figure S1.**
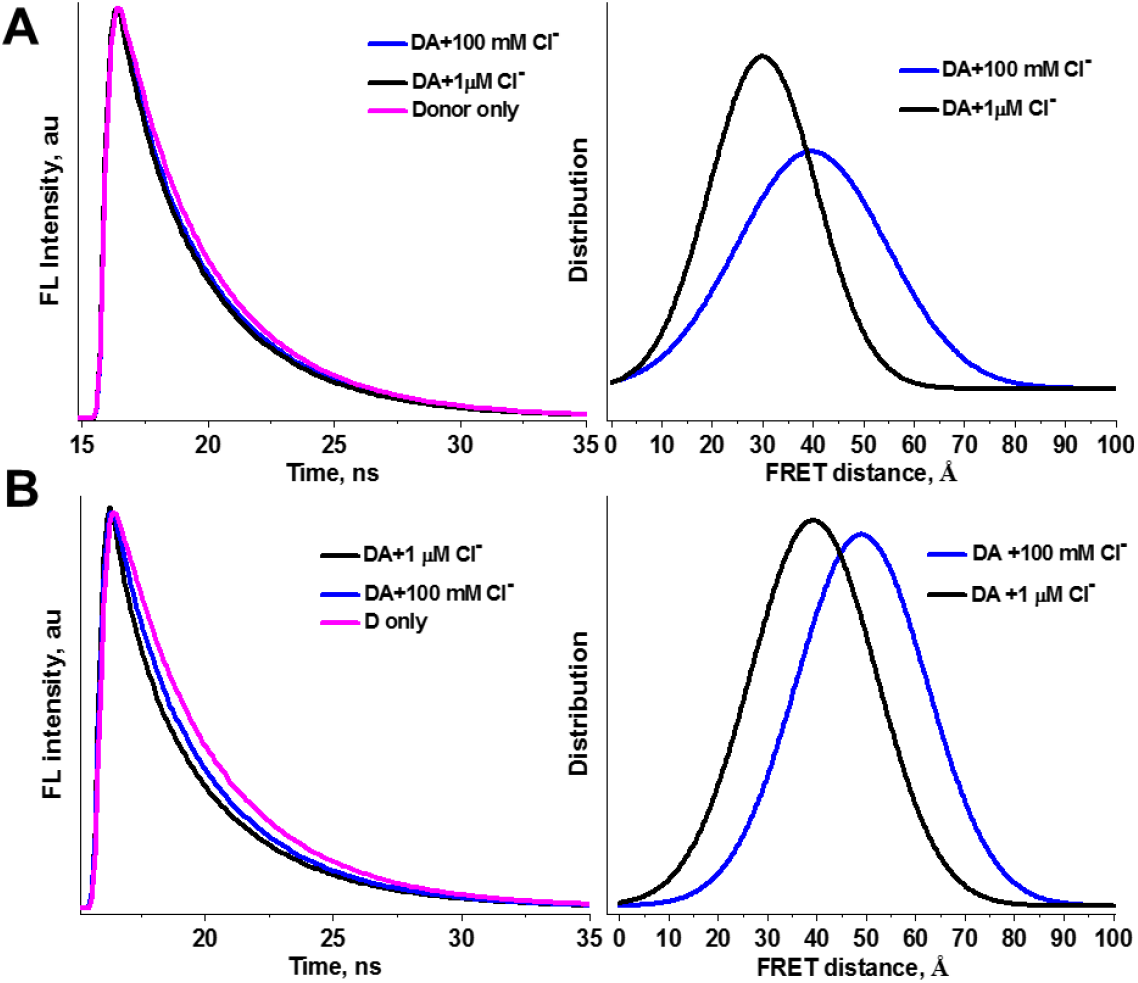
Representative fluorescence decay waveforms and inter-probe FRET distance distributions of the donor labeled and donor-acceptor labeled AC-NTD-RyR2 (A) and BC-NTD-RyR2 (B) constructs in the presence of 0.001 and 100 mM [Cl^−^].

**Figure S2.**
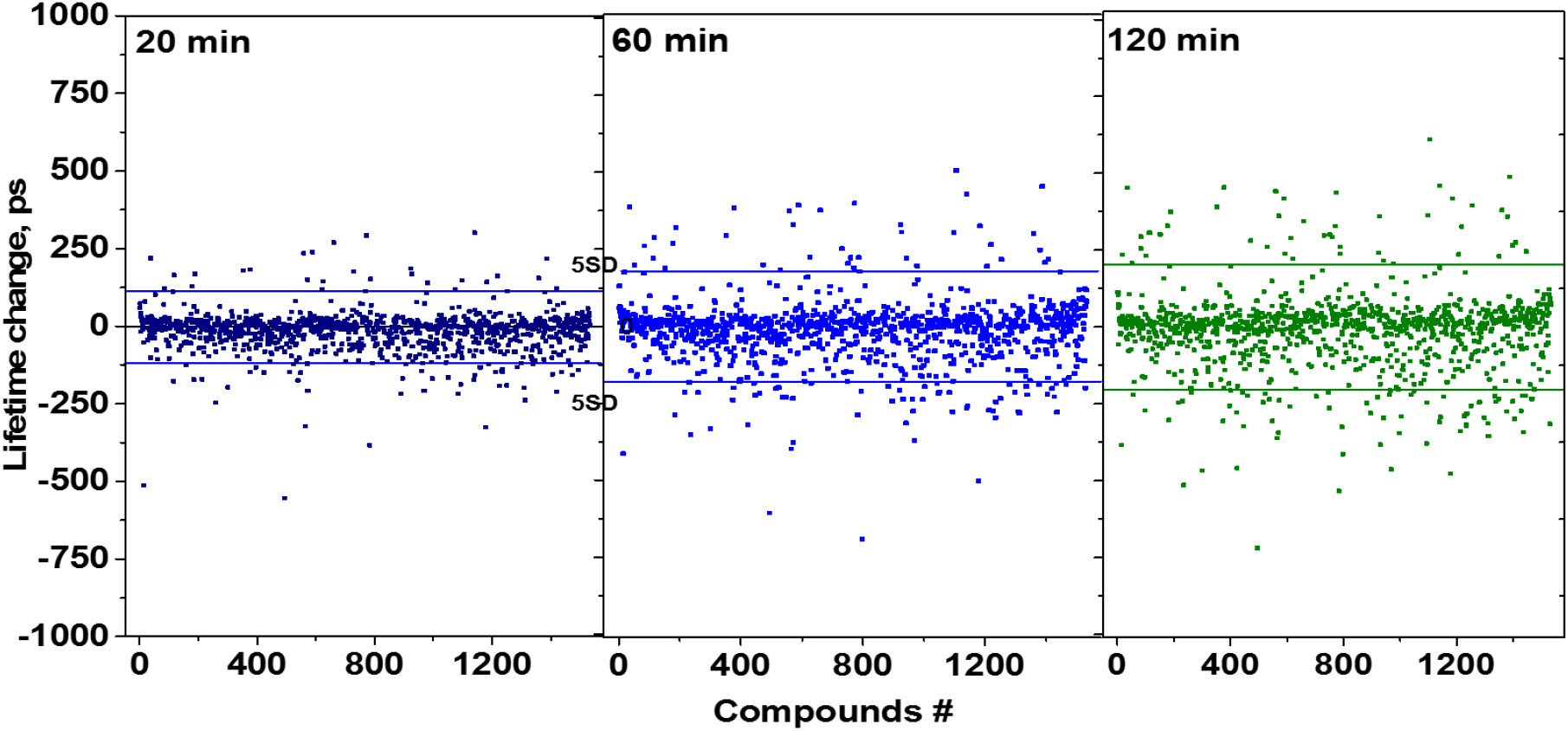
HTS results with AC-NTD-RyR2 biosensor obtained after 20, 60, and 120 min of incubation with 1280 compounds in 1536-well assay plates. Solid lines are ± 5SD in each panel.

**Figure S3.**
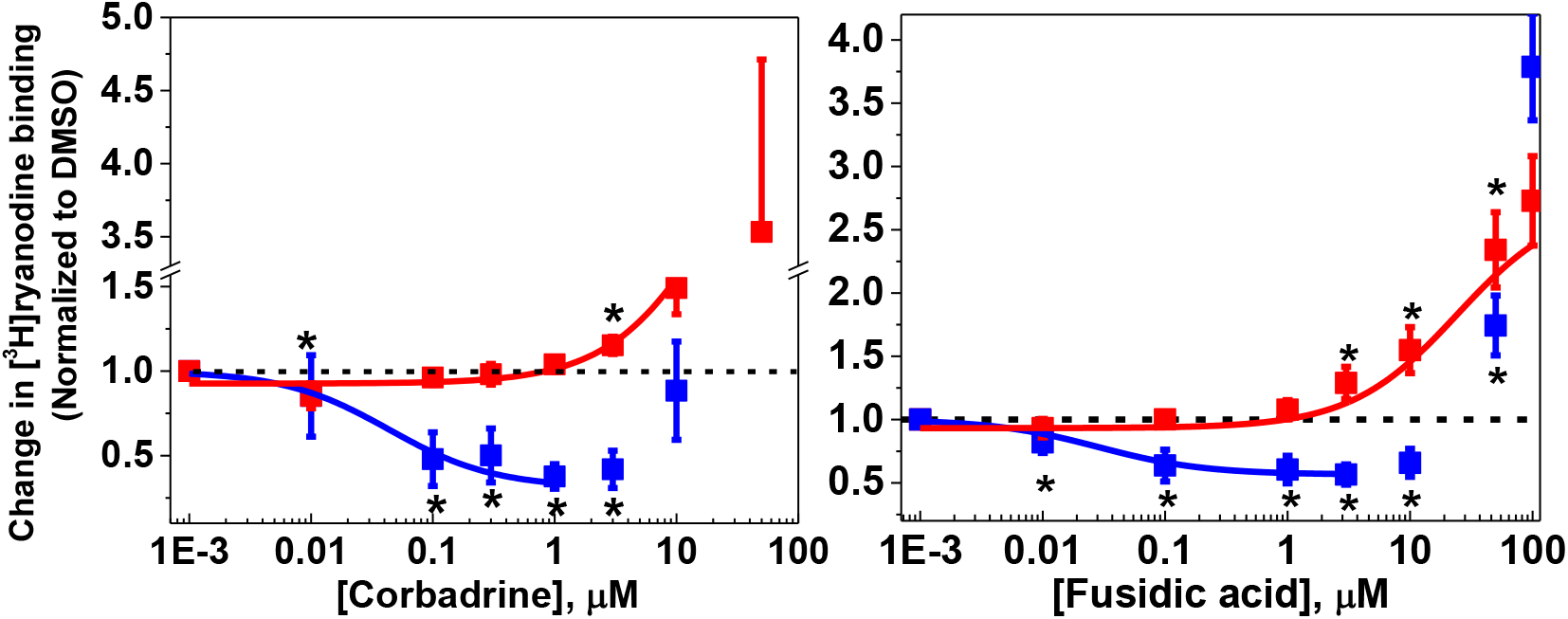
Dose-dependent effect of the corbadrine and fusidic acid on [^3^H]ryanodine binding to porcine skeletal heavy SR (HSR) at 30 nM (blue) and 30 μM (red) free [Ca^2+^]. Results are shown normalized relative to the values for no-drug control (DMSO), respectively, mean ± SE, n = 4. *Significantly different from control using unpaired Student t test, p > 0.05.

**Table S1.** Distance (R) and full-width at half maximum (FWHM) from fitting of FLT-detected intramolecular FRET between donor-acceptor probes in the AC and BC biosensors corresponding to WT- and R420Q-NTR, in the presence of high (100 mM) and low (0.001 mM) [Cl^−^]. SDs were calculated from the average of three experiments.

| [Cl<sup>-</sup>]<br>(mM) | WT BC construct |  | WT AC construct |  | R420Q BC construct |  |
| --- | --- | --- | --- | --- | --- | --- |
|  | R (Å) | FWHM (Å) | R (Å) | FWHM (Å) | R (Å) | FWHM (Å) |
| **0.001** | 40.1±1.0 | 35.2±3.3 | 30.8±0.7 | 25.8±8.3 | 49.6±6.3 | 49.0±7.1 |
| **100** | 51.7±5.4 | 35.7±17.5 | 38.9±1.4 | 37.7±1.9 | 49.5±6.4 | 40.0±21.2 |

**Table S2.** Reproducibility of hits obtained via three HTS runs using the AC- and BC-NTR-RyR2 constructs, after several incubation times with library compounds.

| Incubation<br>time (min) | *n* screens<br>(out of 3) | Number of hits (Hit rate) |  |  |
| --- | --- | --- | --- | --- |
|  |  | AC | BC | AC&BC |
| 20 | 2 | 42 (3.2%) | 9(0.7%) | 4 |
|  | 3 | 13 (1.0%) | 0 |  |
| 60 | 2 | 72 (5.6%) | 30 (2.3%) | 10 |
|  | 3 | 46 (3.5%) | 2 (0.16%) |  |
| 120 | 2 | 63 (4.9%) | 30 (2.3%) | 9 |
|  | 3 | 50 (3.9%) | 4 (0.3%) |  |

**Table S3.**
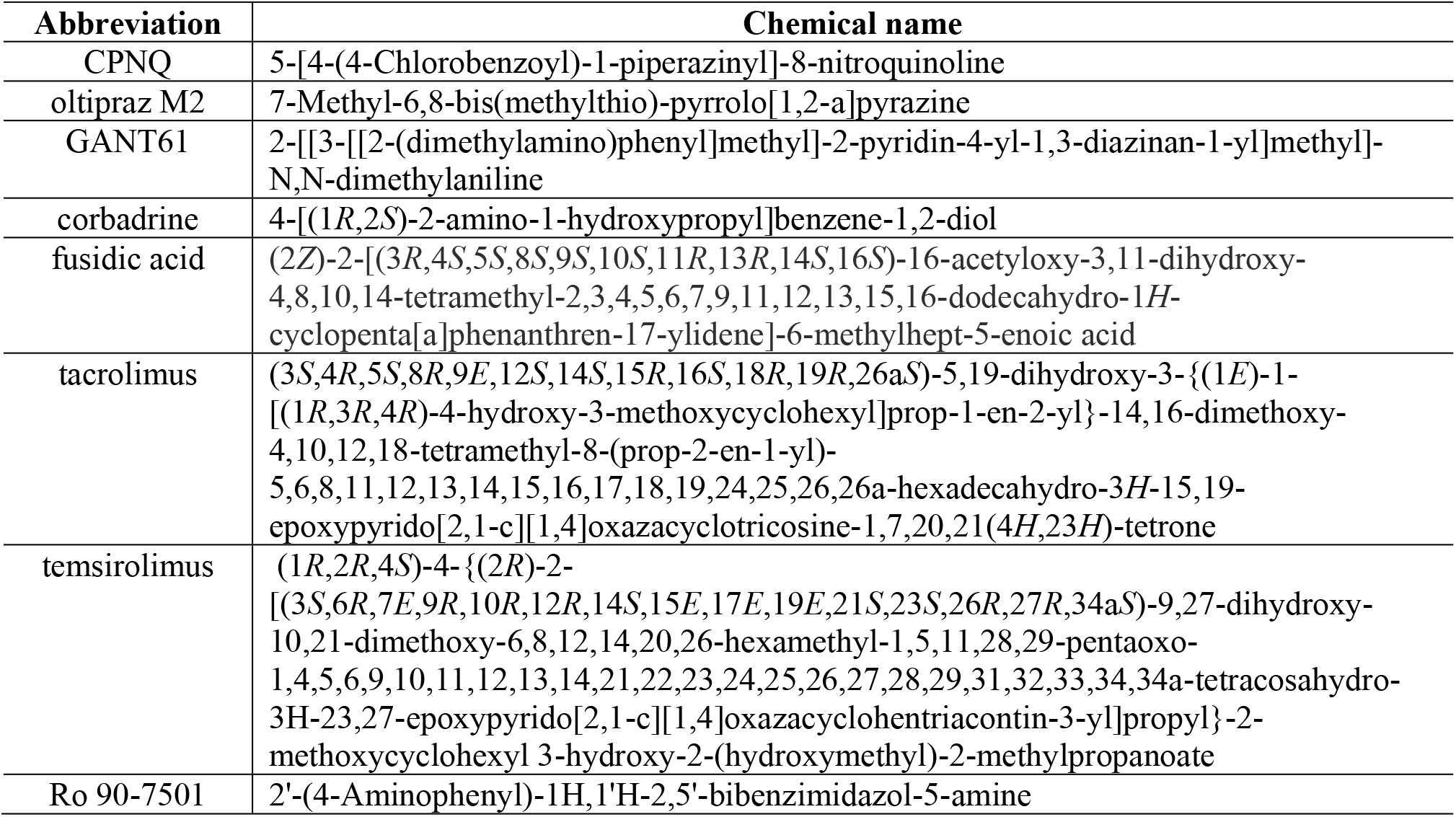
Chemical names and abbreviations of the eight hits studied in this work.

